# A human-specific motif facilitates CARD8 inflammasome activation after HIV-1 infection

**DOI:** 10.1101/2022.10.04.510817

**Authors:** Jessie Kulsuptrakul, Elizabeth A. Turcotte, Michael Emerman, Patrick S. Mitchell

## Abstract

Inflammasomes are cytosolic innate immune complexes that play a critical role in host defense against pathogens but can also contribute to inflammatory pathogenesis. Here, we find that the human inflammasome-forming sensor CARD8 senses HIV-1 infection via site-specific cleavage of the CARD8 N-terminus by the HIV protease (HIV-1^PR^). HIV-1^PR^ cleavage of CARD8 induces pyroptotic cell death and the release of pro-inflammatory cytokines from infected cells, processes that we find are dependent on Toll-like receptor stimulation prior to viral infection. Our evolutionary analyses reveal that the HIV-1^PR^ cleavage site in CARD8 is unique to humans, and that chimpanzee CARD8 does not recognize proteases from HIV or simian immunodeficiency viruses from chimpanzees (SIVcpz). In contrast, SIVcpz does cleave human CARD8, suggesting that SIVcpz was poised to activate the human CARD8 inflammasome prior to its cross-species transmission into humans and implicating the CARD8 inflammasome as a potential driver of HIV pathogenesis.

## Introduction

Simian immunodeficiency viruses (SIVs) are generally non-pathogenic in their natural hosts. In contrast, SIVs often cause disease upon spillover into a new species (e.g., chimpanzees and humans). For instance, human immunodeficiency virus (HIV-1) arose from multiple zoonoses of SIVs from chimpanzees (SIVcpz) and gorillas (SIVgor) ^1^. Similarly, SIVcpz arose from a cross-species transmission event involving the recombination of SIVs from red-capped mangabey (SIVrcm) and mustached monkeys (SIVmus) from Old World monkeys ^2–4^. To varying degrees, SIVs and HIVs in these zoonoses cause chronic immune activation and bystander cell immunopathology that drives progression to acquired immunodeficiency syndrome (AIDS) in the absence of antiretroviral therapy ^5–7^. However, the species-specific interactions driving pathogenesis upon spillover are incompletely understood.

One of the primary selective pressures that shape viral adaptation to a new host, as well as tolerance to persistent infections, is the innate immune system ^8,9^. One class of innate immune sensors form cytosolic immune complexes called inflammasomes, which initiate inflammatory signaling upon pathogen detection or cellular stress ^10^. Inflammasome activation is critical for host defense against a wide range of pathogens; however, auto-activating mutations in inflammasome-forming sensors can also initiate inflammatory pathogenesis that drive autoinflammatory and autoimmune disorders ^11,12^.

The inflammasome-forming sensor caspase recruitment domain-containing protein 8 (CARD8) consists of a disordered N-terminus, a Function-to-Find domain (FIIND), and a caspase activation and recruitment domain (CARD) ^12^. The FIIND, comprised of ZU5 and UPA subdomains, undergoes self-cleavage resulting in two non-covalently associated fragments ^12,13^. Proteasome-dependent degradation of the N-terminus leads to the release and assembly of the C-terminal UPA-CARD, serving as a platform for the recruitment and activation of Caspase-1 (CASP1). Activated CASP1 initiates a lytic, programmed cell death called pyroptosis and the release of pro-inflammatory cytokines including interleukin (IL)-1β and IL-18 ^10,14^. To prevent aberrant release of the UPA-CARD, the dipeptidyl peptidases 8 and 9 (DPP8/9) form an inhibitory complex with CARD8 ^15^. Disruptions to protein homeostasis, including direct (e.g., Val-boroPro) and indirect (e.g., CQ31) inhibition of DPP8/9, cause CARD8 inflammasome activation ^16–18^. These observations have led to the speculation that CARD8 may have evolved to sense cellular stress.

Recently, several examples of pathogen-driven activation of the CARD8 inflammasome have been discovered ^19,20^, including via its recognition of the enzymatic activity of the HIV-1 protease (HIV-1^PR^) ^21^. The ability of HIV-1^PR^ to activate CARD8 was enhanced by enforcing dimerization of the HIV-1^PR^ with a nonnucleoside reverse transcriptase inhibitor (NNRTI). HIV-1^PR^ proteolytic cleavage of the N-terminus of CARD8 causes proteasome-dependent degradation of the CARD8 N-terminal fragment. This ‘functional degradation’ liberates the UPA-CARD fragment for inflammasome assembly and activation. Thus, the N-terminus of CARD8 functions as a ‘tripwire’ to sense and respond to the enzymatic activity of HIV-1^PR^ and other viral proteases ^19,20^. This mechanism is analogous to viral protease sensing by the inflammasome-forming sensor NLRP1 ^22,24,27,28^.

Here, we take an evolution-guided and virological approach to infer the significance of CARD8’s interaction with HIV-1^PR^. We find that human CARD8 has a unique motif among hominoids and Old World monkeys that renders it susceptible to cleavage by HIV-1^PR^ and SIVcpz protease (SIVcpz^PR^), indicating that the precursor viruses to HIV-1 were poised to cleave human CARD8, but do not cleave chimpanzee CARD8. We further demonstrate that human CARD8 can sense HIV-1^PR^ activity and induce inflammasome activation in the context of HIV-1 infection *in vitro*, but only if the target cells are first activated (e.g., via Toll-like receptor (TLR) priming) prior to viral challenge. CARD8 activation by HIV-1 infection leads to cell death and IL-1β secretion, thereby suggesting a model for a human-specific pathogenic response to lentiviral infection.

## Results

### A human-specific motif allows CARD8 to detect protease activity from multiple HIV strains

The HIV-1 protease (HIV-1^PR^) cleaves human CARD8 between phenylalanine (F) 59 (P1) and F60 (P1’) (**Figure 1A**) ^21^. While the amino acid P1 site, F59, is invariant among hominoids, gibbons, and Old World monkeys, only human CARD8 has a phenylalanine at the P1’ site, F60 (**Figure 1A**). We first established the conditions required for HIV^PR^ cleavage of CARD8 by assessing the ability of two HIV-1 group M proviruses (HIV-1_LAI_ subtype B and HIV-1_Q23_ subtype A) and HIV-2_ROD_ to cleave human CARD8. Indeed, we found that wildtype (WT) human CARD8 with an N-terminal mCherry fusion is cleaved upon overexpression of HIV-1 Gag-Pol, resulting in a ∼33 kDa product (**Figure 1B** and **1C**, top blot). Cleavage of CARD8 by Gag-Pol was dependent on HIV-1^PR^ activity as the HIV^PR^ inhibitor Iopinavir (LPV) blocked both Gag processing of p55^gag^ to p41^gag^ and p24^gag^ and CARD8 cleavage (**Figure 1C**, top and middle blot). Moreover, we found that human CARD8 is also susceptible to HIV-2_ROD_ protease activity (**Figure 1C**) indicating that cleavage of human CARD8 is conserved across HIV-1 and HIV-2. To assess the significance of the amino acid variation at the F60 P1’ site of CARD8, we next replaced human CARD8 F60 (WT) with either a leucine (L; found in chimpanzee, bonobo, and gorilla) or a serine (S; found in gibbons and Old World monkeys) (**Figure 1A**). HIV^PR^ cleavage of WT human CARD8 (F60) was much more efficient than cleavage of human CARD8 F60L or F60S (**Figure 1D**), consistent with prior findings that an alanine at position 60 also blocks HIV^PR^ ^21^. These results indicate that species-specific variation at position 60 impacts CARD8 recognition of HIV^PR^ activity.

**Figure 1:**
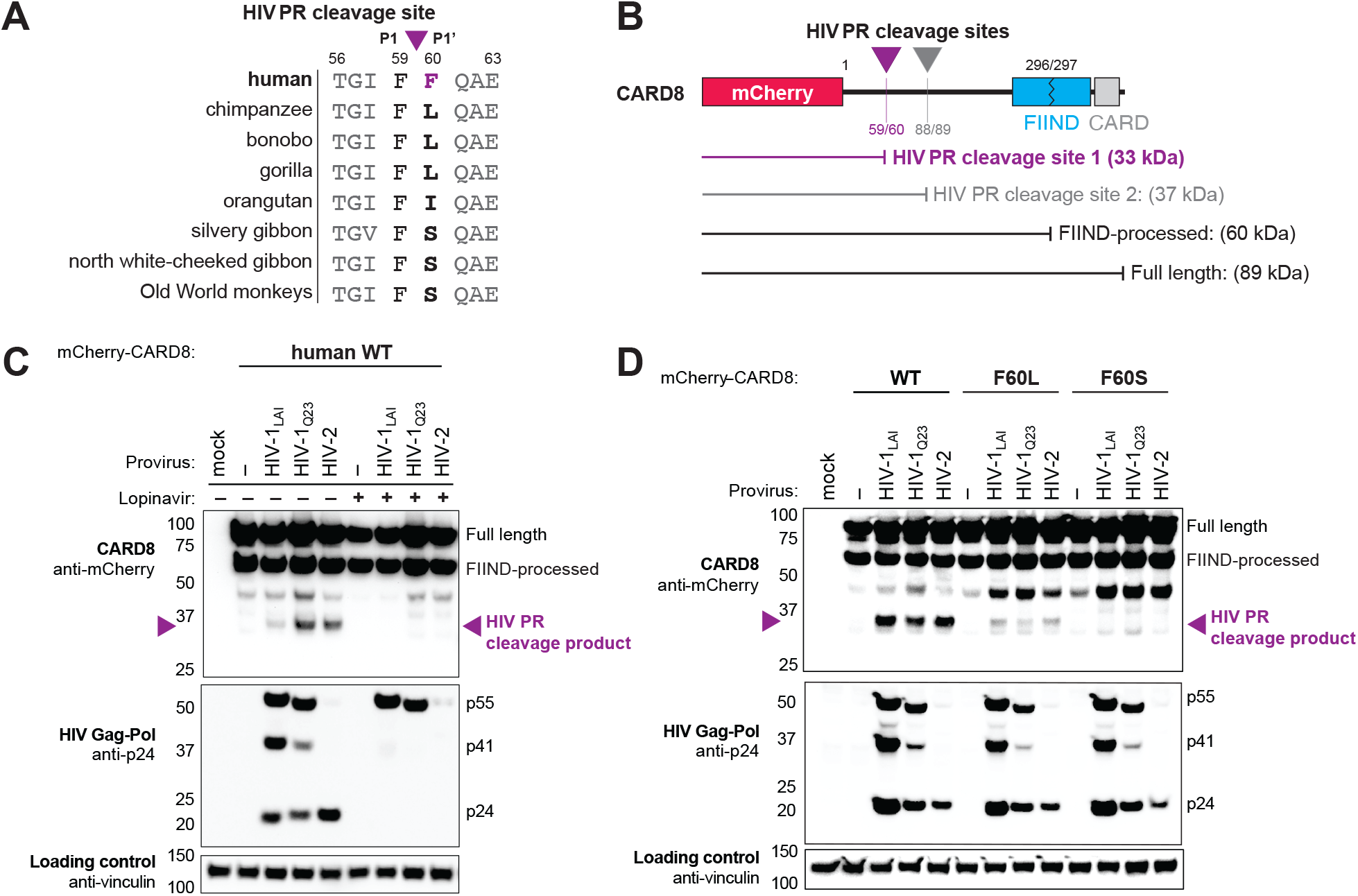
A human-specific motif allows CARD8 to detect protease activity from multiple HIV strains. **A)** Alignment of primate CARD8 protein sequences. The HIV protease (HIV PR) cleavage site is indicated by a purple triangle between F59 (P1) and F60 (P1’). Numbering is based on human CARD8. **B)** Depiction of the mCherry-CARD8 used in cleavage assays in C) and D). The predicted molecular weights (kDa) for full length, FIIND-processed, or HIV PR cleavage products are indicated. FIIND, function-to-find domain; CARD, caspase activation and recruitment domain. **C)** HEK293T cells were transfected with a construct encoding N-terminally mCherry-tagged wildtype (WT) CARD8 and indicated HIV proviral constructs in the presence (‘+’) or absence (‘–’) of 10 μM lopinavir, an HIV PR inhibitor. Immunoblotting was carried out for CARD8 cleavage, HIV PR activity, and vinculin (loading control) as indicated. The band at ∼45kDa is the result of cleavage by the 20S proteasome ^23^. **D)** HEK293T cells were transfected with a construct encoding N-terminally mCherry-tagged WT, F60L, or F60S CARD8 and indicated HIV proviral constructs. Immunoblotting was performed as in C).

### Natural variation in CARD8 alters sensing of SIVcpz protease activity

We next asked if HIV^PR^ cleavage of CARD8 was an ancestral function of SIVcpz or if that functionality instead emerged following cross-species transmission and adaptation to humans. SIVcpz_EK505_ and SIVcpz_LB7_ represent lineages that gave rise to HIV-1 group N and M viruses, respectively ^1,25,26^. Like HIV-1 and HIV-2 proteases, we found that both SIVcpz proteases (SIVcpz^PR^) cleaved human CARD8 (**Figure 2A**), suggesting that SIVcpz^PR^ had a pre-existing ability to cleave human CARD8 prior to spillover. To deduce whether or not this cleavage is unique to humans, we also tested SIVcpz^PR^ ability to cleave chimpanzee CARD8 (**Figure 2A**) and F60L and F60S human CARD8 variants (**Figure 2B**) and found that none of the other CARD8 variants could be cleaved by SIVcpz^PR^. These data suggest that SIVcpz^PR^ was poised to cleave human CARD8 prior to its zoonosis to humans, and human CARD8 is uniquely susceptible to HIV and SIVcpz protease cleavage.

**Figure 2:**
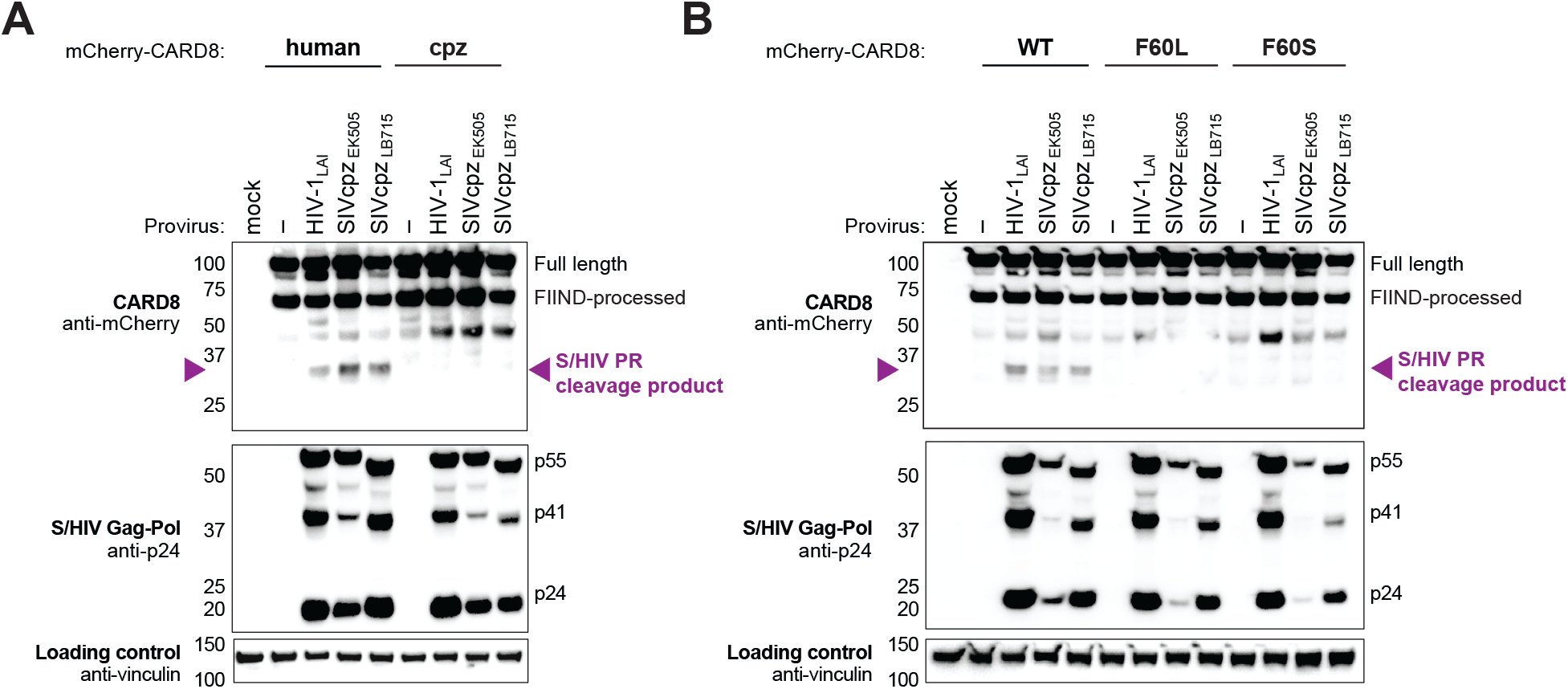
Natural variation in CARD8 alters sensing of SIVcpz^PRx^ activity. **A)** HEK293T cells were transfected with a construct encoding N-terminally mCherry-tagged human or chimpanzee (cpz) CARD8 and indicated provirus constructs. Immunoblotting was carried out for CARD8 cleavage, HIV/SIV protease (S/HIV PR) activity, and vinculin (loading control) as indicated. The S/HIV PR cleavage product is indicated by a purple triangle. FIIND, function-to-find domain. **B)** HEK293T cells were transfected with a construct encoding N-terminally mCherry-tagged wildtype (WT), F60L, or F60S CARD8 and indicated proviral constructs. Immunoblotting was performed as in A).

### HIV-1 infection activates the inflammasome in primed THP-1 cells

We next sought to determine the significance of CARD8 cleavage and activation in the context of HIV-1 infection. Treatment with nonnucleoside reverse transcriptase inhibitors (NNRTIs) induces premature Gag-Pol dimerization and HIV-1^PR^ activity ^29,30^, which was previously shown to be required for CARD8 activation in HIV-1 infected cells ^21^. However, other studies have shown that Gag-Pol can reach intracellular concentrations that are sufficient to dimerize and activate HIV-1^PR^ during HIV-1 infection ^31^. Thus, to determine if CARD8 inflammasome activation can occur during HIV-1 infection in the absence of small molecule-induced HIV-1^PR^ dimers, we infected the human leukemia monocytic cell line THP-1 with either HIV-1_LAI_ or VSV-g pseudotyped HIV-1_LAI_ (HIV-1_LAI-VSVG_) at a multiplicity of infection <1 in the absence of NNRTIs. As a positive control for inflammasome activity, uninfected cells were also treated with VbP, which specifically activates the CARD8 inflammasomes in THP-1 cells ^16^. For both HIV-1 infected and VbP-treated THP-1 cells, we observed an increase in cell death compared to mock infected controls as measured by uptake of the membrane impermeable dye propidium iodide (PI) (**Figure 3A**). In parallel, we evaluated if HIV-1 infection also results in the release of IL-1β. Consistent with prior reports, neither HIV-1 infection nor VbP alone led to an increase in IL-1β levels (**Figure 3B**) ^32–34^. We reasoned that the lack of cytokine production may either be an intrinsic property of CARD8 ^34^, or alternatively, require a signal 1 (e.g., a TLR agonist) to transcriptionally upregulate or ‘prime’ IL-1β and/or inflammasome components ^35^. Thus, we assessed inflammasome activation by HIV-1 infection or VbP treatment with and without priming of THP-1 cells using the TLR2 agonist Pam3CysSerLys4 (Pam3CSK4). We found that HIV-1_LAI_ and HIV-1_LAI-VSVG_ infection induces cell death in both primed and unprimed cells; however, priming significantly increased cell death upon HIV-1 infection (**Figure 3A**). In contrast, release of IL-1β after infection with HIV-1 was entirely dependent on TLR priming (**Figure 3B**). Additionally, we observed Gag processing of p55^gag^ to p41^gag^ and p24^gag^ in cytoplasmic lysates of HIV-1 infected cells, consistent with prior studies demonstrating that HIV-1^PR^ is active in the cytoplasm (**Figure 3C**) ^31,36^. Thus, HIV-1 infection activates an inflammasome response in THP-1 cells, which requires priming for IL-1β secretion but not cell death.

**Figure 3:**
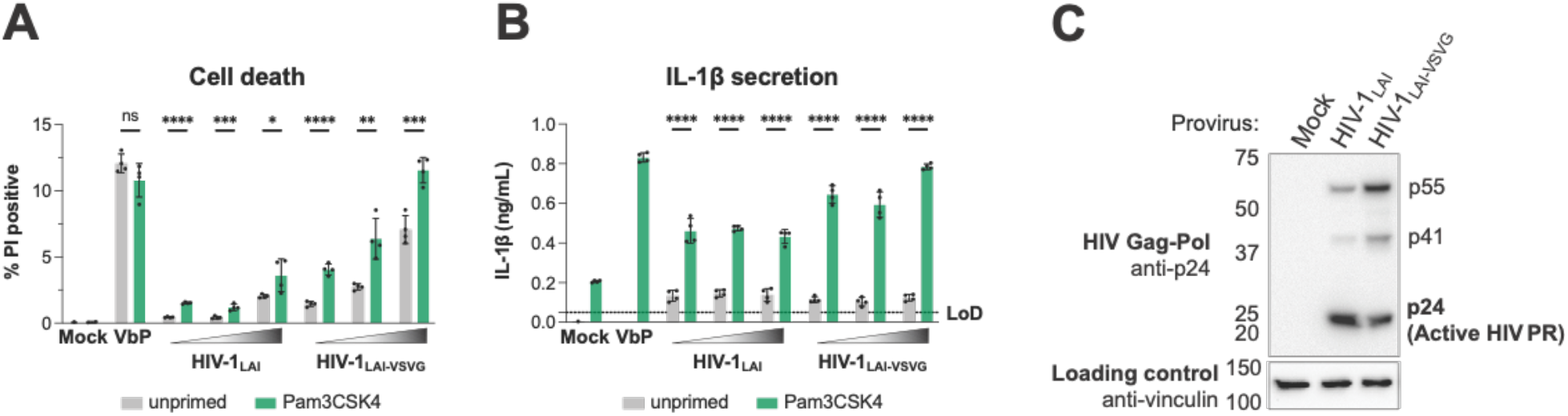
HIV-1 infection activates the inflammasome in THP-1 cells. **A-B)** THP-1 cells were either left unprimed or primed with TLR2 agonist Pam3CSK4 4-6 hours before treatment with 10 μM VbP or different multiplicities of infection (MOIs) for HIV-1_LAI_ or HIV-1_LAI-VSVG_. HIV-1 MOIs (increasing from left to right) had approximately 4%, 8%, and 45% HIV-1_LAI_-infected cells and 33%, 53%, and 90% HIV-1_LAI-VSVG_-infected cells, as determined by flow cytometry of intracellular p24^gag^ of unprimed cells 24 hours post-infection. Inflammasome responses were measured 24 hours following VbP treatment or HIV-1 infection. **A)** Cell death is reported as the percent of propidium iodide (PI) positive cells. **B)** IL-1β levels were measured using the IL-1R reporter assay. The dotted line indicates limit of detection (LoD). Datasets represent mean ± SD (n = 4 biological replicates). *P* values were determined by unpaired two-sided t-tests using GraphPad Prism 9. ns= not significant, p< 0.05, * *p< 0.01, * * *p< 0.001, * * * *p< 0.0001. **C)** THP-1 cells were mock infected or infected with HIV-1_LAI_ or HIV-1_LAI-VSVG_ using the middle HIV dose from A-B), yielding 8% and 53% infection, respectively. Immunoblotting using cytoplasmic lysates was carried out for HIV protease (HIV PR) activity, and vinculin (loading control) as indicated 24 hours post-infection.

### HIV-1 inflammasome activation is dependent on a human-specific motif in CARD8

To determine if inflammasome activation upon HIV-1 infection is dependent on the human-specific motif in CARD8, we first generated clonal THP-1 *CARD8* knockout (KO) cells via CRISPR/Cas9. We confirmed the absence of full length (∼62kDa) and FIIND-processed (∼29kDa) CARD8 in THP-1 *CARD8* KO cell lines by immunoblotting with an antibody specific to the CARD8 C-terminus (**Figure 4A**). To functionally test the THP-1 *CARD8* KO cell lines, we primed WT or *CARD8* KO THP-1 cells with Pam3CSK4 then treated with either VbP, which activates the CARD8 inflammasome, or the ionophore nigericin, which specifically activates the NLRP3 inflammasome, and measured cell death and IL-1β secretion. As expected, WT but not *CARD8* KO THP-1 cells responded to VbP, whereas both cell lines underwent cell death and IL-1β secretion in response to nigericin, indicating that the *CARD8* KO THP-1 cells retained responsiveness to other inflammasome agonists (**Figure 4B**). Similar to our observations with VbP, we found that IL-1β secretion and cell death were significantly reduced in Pam3CSK4-primed, HIV-1_LAI_ or HIV-1_LAI-VSVG_ infected *CARD8* KO versus WT THP-1 cells (**Figure 4C**). These results indicate that HIV-1-induced inflammasome activation in THP-1 cells is dependent on CARD8.

**Figure 4:**
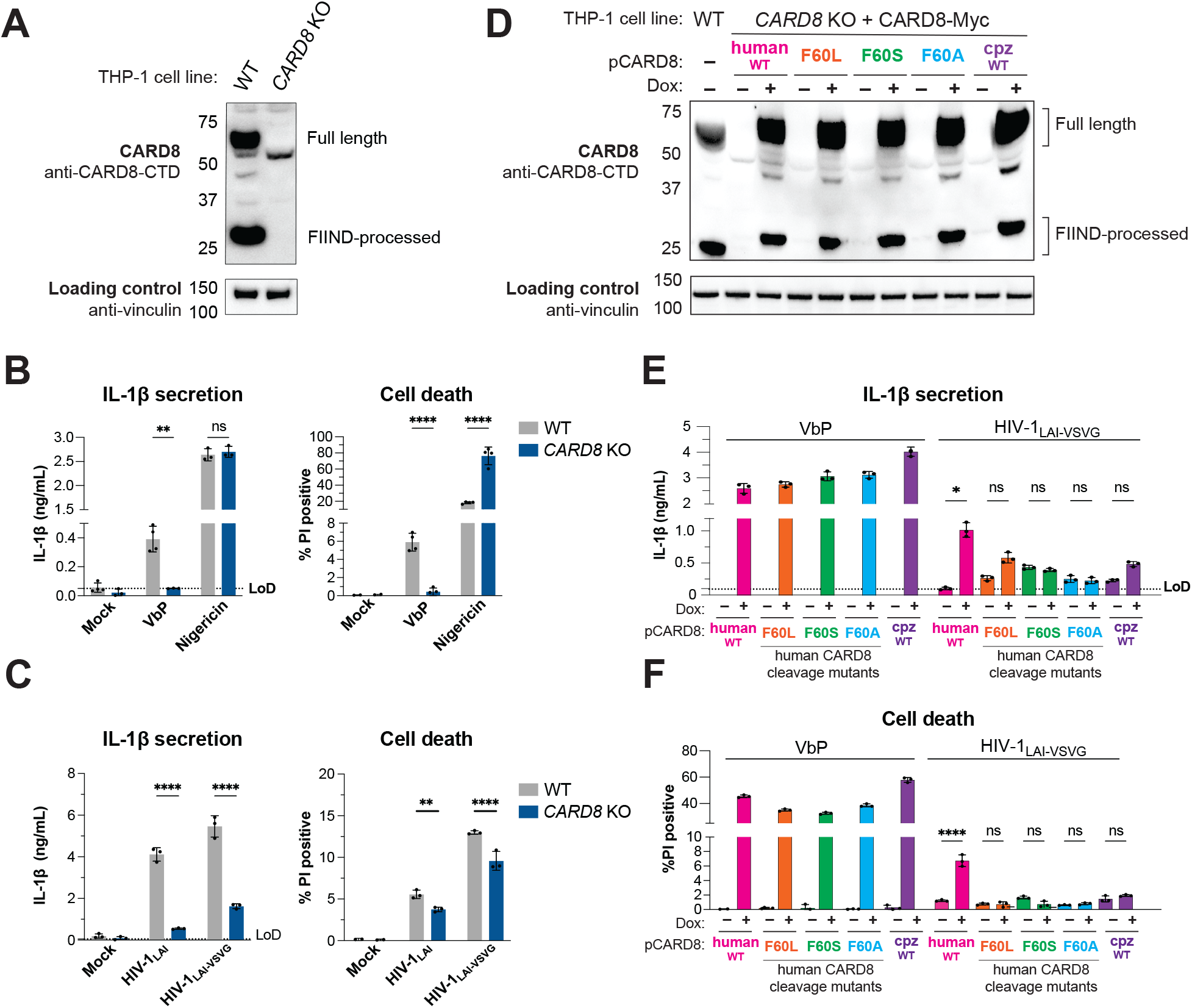
HIV-1 inflammasome activation is dependent on a human-specific motif in CARD8. **A)** Immunoblot of wildtype (WT) versus *CARD8* knockout (KO)THP-1 cells was carried out for CARD8 expression using endogenous antibody against CARD8 C-terminal domain (CTD) and loading control (vinculin). FIIND, function-to-find domain. **B)** WT or *CARD8* KO THP-1s were primed for 4-6 hours with TLR2 agonist Pam3CSK4 and either treated with mock, 10 μM VbP, or 5 μg/mL nigericin, and probed for (*left*) supernatant IL-1β using an IL-1R reporter assay and (*right*) cell death via percent of propidium iodide (PI) positive cells. Treatment for VbP and nigericin were for 24 and 2 hours, respectively. Dotted line indicates limit of detection (LOD). Datasets represent mean ± SD (n = 3 biological replicates). *P* values were determined by unpaired two-sided t-tests using GraphPad Prism 9. ns=not significant, *p< 0.05, * *p< 0.01, * * *p< 0.001, * * * *p< 0.0001. **C)** WT or *CARD8* KO THP-1s were primed for 4-6 hours with Pam3CSK4 and either treated with mock, HIV-1_LAI_ or HIV-1_LAI-VSVG_ and probed 24 hours post-infection for (*left*) IL-1β levels and (*right*) cell death as in B). HIV-1_LAI_ and HIV-1_LAI-VSVG_ infections were done at a multiplicity of infection (MOI) <1 such that only 30-40% of WT cells were infected as determined by intracellular p24^gag^. **D)** *CARD8* KO THP-1 lines complemented with different doxycycline (dox)-inducible CARD8 variants (pCARD8) were left uninduced or induced for 18 hours. Immunoblot of wildtype (WT) or complemented *CARD8* KO THP-1 lines treated with (‘+’) or without (‘–’) dox was carried out for CARD8 expression as described in A). **E-F)** Complemented *CARD8* KO lines were left uninduced or dox-induced as described in D) then primed for 4-6 hours with Pam3CSK4 and treated with either 10 μM VbP or HIV-1_LAI-VSVG_ then assessed for **E)** IL-1β secretion and **F)** cell death as previously described in B) and C), respectively. HIV-1_LAI-VSVG_ infection was done at the same MOI used in C).

Finally, to determine if CARD8-dependent inflammasome activation by HIV-1 infection requires HIV-1^PR^ cleavage of CARD8, we used a doxycycline inducible system to complement *CARD8* KO THP-1 cells with either WT CARD8 or CARD8 cleavage mutants (**Figure 4D**) and probed for subsequent inflammasome activation. We found that *CARD8* KO THP-1 cells complemented with WT CARD8 underwent IL-1β secretion and cell death in response to both VbP and HIV-1 infection in a doxycycline and Pam3CSK4 dependent manner (**Figure 4E and 4F**), indicating that HIV inflammasome activation is CARD8-dependent. In order to test if HIV-1^PR^ cleavage of CARD8 is required for inflammasome activation by HIV infection, in parallel, we complemented *CARD8* KO THP-1 cells with the CARD8 cleavage mutants F60L, F60S, F60A, or full-length chimpanzee CARD8. All complemented *CARD8* KO THP-1 cells underwent IL-1β secretion and cell death in response to VbP in doxycycline-treated cells, demonstrating functional CARD8 expression (**Figure 4E and 4F, left sides**). In contrast, HIV-1 infection only induced IL-1β secretion and cell death in *CARD8* KO THP-1 lines that were complemented with WT human CARD8 (**Figure 4E and 4F, right sides**). Thus, HIV-1 infection induces inflammasome activation via CARD8 cleavage by HIV^PR^ in a manner that depends on a human-specific motif at the site of cleavage.

## Discussion

HIV-1 arose from the cross-species transmission of SIVcpz from chimpanzees. Here, we considered the evolutionary context of CARD8-lentivirus protease interactions. We found that a substitution that arose in humans during CARD8 evolution renders it susceptible to cleavage and activation by proteases from both HIV and the HIV-1 precursor SIVcpz, implying that SIVcpz^PR^ was poised to cleave human CARD8 prior to spillover of SIVcpz into humans and did not occur as a result of adaptation to the human host. We also found that SIVcpz cannot cleave chimpanzee CARD8, suggesting that CARD8 sensing of lentiviral proteases may uniquely play a role in HIV pathogenesis.

The F59-F60 motif that confers human CARD8 with the unique capacity to sense HIV/SIVcpz^PR^ is conserved across all humans, suggesting a genetic sweep occurred in favor of a phenylalanine at position 60. HIV-1 emerged within the past century ^4^ and therefore could not have driven the evolution of the HIV-1^PR^ cleavage site in human CARD8. In contrast, human *CARD8* is highly polymorphic, and multiple regions of the N-terminus of CARD8 show strong evidence of positive selection ^20^. Moreover, the HIV-1^PR^ cleavage site in CARD8 overlaps with a site that is cleaved by the coronavirus 3CL protease ^20^. Thus, although it is possible that the human-specific F60 was fixed stochastically or as a passenger mutation, we favor a scenario in which human CARD8 sensing of HIV-1^PR^ arose as a consequence of CARD8 adaptation to another virus ^19,20^. This hypothesis is consistent with SIV-infected primates exhibiting reduced ^5^ or no overt pathogenesis ^37^ relative to HIV-infected patients who quickly progress to AIDS without treatment ^1^.

HIV-1 disease progression to AIDS is characterized by dramatic depletion of CD4 T cells including via pyroptosis ^38^ and chronic inflammation accompanied by high levels of plasma cytokines including IL-1 ^39,40^. As such, multiple inflammasomes have previously been implicated for HIV-dependent inflammasome activation, although the exact mechanisms have remained unclear ^41,42^. Here, we show that HIV infection induces a CARD8-dependent inflammasome response resulting in cell death and IL-1β secretion by recognizing HIV^PR^ activity. Our findings suggest a role for the CARD8 inflammasome in HIV-1 pathogenesis and underscore the importance of innate immune detection of virus proteolytic activity during viral infection by innate immune sensors like CARD8.

Interestingly, we find that inflammasome responses downstream of CARD8 are modulated by TLR stimulation (**Figure 3**). CARD8-dependent cell death is enhanced by but can occur independent of TLR priming, suggesting that TLR stimulation may modulate CARD8-dependent cell death or permit the engagement of other cell death sensors and/or pathways during HIV-1 infection ^41,42^. On the other hand, IL-1β secretion following HIV-1 infection is strictly dependent on priming, offering a potential explanation for conflicting reports as to whether or not primary CD4 T cells undergo pyroptosis and induce IL-1β secretion in response to HIV-1 infection ^38,43^. HIV-1 pathogen-associated molecular patterns (i.e., viral nucleic acids) and/or circulating microbial ligands from gut epithelial breakdown, a hallmark of acute HIV-1 disease ^44^, are potential sources for priming of HIV-1 target cells *in vivo*. Future studies are required to uncover the mechanisms by which priming modulates CARD8 inflammasome activation.

Our findings also demonstrate that inflammasome priming is crucial for eliciting a full CARD8 inflammasome response, and that HIV-1 infection at low multiplicity of infection can occur in the absence of NNRTIs. In contrast, Wang et al. discovered that HIV^PR^ is sensed by CARD8 using NNRTIs to enforce HIV^PR^ cytosolic activity as a means to clear latently HIV-infected cells ^21^. We speculate that therapeutic strategies that leverage HIV^PR^-dependent CARD8 inflammasome activation may be bolstered by adjuvants that induce TLR signaling ^21,45,46^. Our finding that HIV-1 infection is sufficient to induce inflammasome activation, along with the presence of CARD8 in relevant T cell populations ^16,33^, also suggests that CARD8 may contribute to HIV pathogenesis. For instance, IL-1β induces the differentiation of Th17 cells ^47^, a highly HIV-susceptible CD4 T cell subtype, and recruits other target immune cells ^48^. Additionally, CARD8 inflammasome activation may contribute to CD4 T cell depletion via CARD8-dependent pyroptosis or lead to increased pathogenic inflammatory responses ^49^.

Taken together, our work highlights how even minor, single amino acid changes can have dramatic, species-specific impacts on innate immune sensing and pathogenesis, and provides a model to explain, in part, the unique susceptibility of humans to HIV pathogenesis.

## Acknowledgements

We thank everyone in the Emerman and Mitchell labs, Amandine Chantharath for assistance with cloning, Matt Daugherty, Emily Hsieh, and Brian Tsu for critical reading of the manuscript, Janet Young for helpful discussions, Melissa Kane of the University of Pittsburgh for kindly providing the doxycycline inducible plasmid (pLKO-puro) used for complementation experiments, and the Fred Hutchinson Shared Resources Genomics, Flow Cytometry, and Specimen Processing & Research Cell Bank cores. Lopinavir (LPV) and p24^gag^ antibody (ARP-3537) were provided by the AIDS Reagent Program, Division of AIDS, NIAID, NIH. This work was supported by grants from the National Institutes of Health (NIH) (DP2 AI 154432-01) and the Mallinckrodt Foundation to P.S.M; NIH grants DP1 DA051110-03 to M.E., and University of Washington Cellular and Molecular Biology Training Grant (T32 GM007270) to JK.

## Author Contributions

Conceptualization: JK, ME, PSM; Methodology and investigation: all authors; Visualization: JK, ME, PSM; Supervision: ME, PSM; Writing – original draft: JK, PSM, ME; Writing – review & editing: All authors

## Declaration of Interests

The authors declare no competing interests.

## Methods

### Plasmids

psPAX2 and pMD2.G were gifts from Didier Trono (Addgene). The dox-inducible pLKO-puro vector ^50^ was a gift from Melissa Kane. Infectious molecular clones for SIVcpz_EK505_ and SIVcpz_LB715_ were gifts from Beatrice Hahn ^25,26^. HIV-1_Q23_ Δenv provirus was a gift from Julie Overbaugh ^51^. HIV-1_LAI_ and HIV-2_Rod_ were previously described ^52,53^. For CARD8 cleavage assays, the coding sequences of human CARD8 (NCBI accession NP_001171829.1) and chimpanzee CARD8 (NCBI accession XM_024351500.1) were cloned into the pcDNA3.1 backbone (Addgene) with an N-terminal mCherry tag using BamHI and EcoRI cut sites. For dox-inducible complementation assays, the coding sequences of human and chimpanzee CARD8 were cloned into the pLKO-puro backbone using the Sfil site. Point mutations were introduced using overlapping PCR. Full list of primer sequences can be found in **Table S1**.

### Cell Culture

THP-1 cells (ATCC) were cultured in RPMI (Invitrogen) with 10% FBS, 1% penicillin/streptomycin antibiotics, 10 mM HEPES, 0.11 g/L sodium pyruvate, 4.5 g/L D-Glucose and 1% Glutamax. HEK 293T (ATCC) were cultured in DMEM (Invitrogen) with 10% FBS and 1% penicillin/streptomycin antibiotics. All puromycin selections were done at 0.5 μg/mL. For complemented dox-inducible lines, tetracycline-free FBS (Sigma) was used to prevent background CARD8 expression. All lines routinely tested negative for mycoplasma bacteria (Fred Hutch Specimen Processing & Research Cell Bank).

### Immunoblotting

Cells were washed once with 1xPBS before harvesting in NP-40 buffer with protease inhibitor (200 mM NaCl, 50 mM Tris pH 7.4, 0.5% NP-40 Alternative, 1 mM dithiothreitol, and Roche Complete Mini, EDTA-free tablets; catalog no. 11836170001). Cytoplasmic lysates were clarified via centrifugation and combined with 4x NuPage LDS sample Buffer (Invitrogen) containing 10% β-mercaptoethanol and boiled for 5-10 minutes. Samples were run on a 4-12% SDS-PAGE gel using morpholineethanesulfonic acid (MES) buffer, transferred to a nitrocellulose membrane using a Pierce G2 Fast Blotter (Thermo Scientific), blocked in 5% nonfat milk then probed for with primary antibodies diluted in 1% milk for mCherry (for CARD8 cleavage), p24^gag^ (for HIV^PR^ activity), CARD8 C-terminus (for knockout validation and complementation), and vinculin (loading control). Blots were washed three times with PBS-T (0.1% Tween-20), incubated with secondary HRP-conjugated antibodies, washed three times again, and then developed with SuperSignal West Femto Maximum Sensitivity Substrate (Fisher Scientific). Further antibody specifications and clone info are described in **Table S2**.

### CARD8 cleavage assay

HEK293T cells were seeded at 1.5-2 × 10^5^ cells/well in 24-well plates the day before transfection using TransIT-LT1 reagent at 1.5μL transfection reagent/well (Mirus Bio LLC). 100 ng of indicated constructs encoding a N-terminal mCherry tagged CARD8 were co-transfected into HEK293T cells with either 400 ng of pcDNA3.1 empty vector (‘–‘), or 400 ng of HIV provirus or SIVcpz provirus. HIV Δenv proviruses were used for immunoblots in **Figure 1**, while infectious HIV and SIVcpz provirus were used for immunoblots in **Figure 2**. Cytoplasmic lysates were harvested 24 hours post-transfection and immunoblotted as described above.

### *CARD8* knockout generation

*CARD8* knockout THP-1 cells were generated similarly to *NLRP1* knockouts described previously ^28^. Briefly, a *CARD8* specific sgRNA was designed using CHOPCHOP ^54^, and cloned into a plasmid containing U6-sgRNA-CMV-mCherry-T2A-Cas9 using ligation-independent cloning. THP-1 cells were electroporated using the BioRad GenePulser Xcell. After 24 hours, mCherry-positive cells were sorted and plated for cloning by limiting dilution. Monoclonal lines were validated as knockouts by deep sequencing and OutKnocker analysis, as described previously ^55,56^. Knockout lines were further validated by immunoblot and functional assays. sgRNA used to generate knockouts are described in **Table S1**.

### CARD8 complementation

HEK293T were seeded at 2 × 10^5^ cells/well in 6-well plates the day before transfection using TransIT-LT1 reagent (Mirus Bio LLC) at 5.8 μL transfection reagent/well. Cells were co-transfected with pLKO-CARD8, psPAX2, and pMD2.G and media was replaced the next day. Virus was harvested two days post-transfection and underwent one freeze thaw cycle at −80°C before transducing *CARD8* KO THP-1 cells. *CARD8* KO THP-1 cells were seeded at 2 × 10^5^ cells/well in 6-well plates and transduced with 800 μL virus in the presence of 1 μg/mL polybrene via spinoculation at 1100 x *g* for 30 minutes at 30°C then puro-selected 24 hours post-transduction.

### HIV-1_LAI_ and HIV-1_LAI-VSVG_ production

293T cells were seeded at 2 × 10^5^ cells/well in 6-well plates the day before transfection using TransIT-LT1 reagent (Mirus Bio LLC) at 3 μL transfection reagent/well as previously described ^57^. For HIV-1 production, 293Ts were transfected with either 1 μg/well HIV_LAI_ proviral DNA or 1 μg/well HIV_LAI_ Δenv DNA and 500 ng/well pMD2.G for HIV-1_LAI_ and HIV-1_LAI-VSVG_, respectively. One day post-transfection, media was replaced. Two or three days post-transfection, viral supernatants were collected and filtered through a 20 μm filter and aliquots were frozen at - 80°C. HIV-1_LAI_ and HIV-1_LAI-VSVG_ proviruses were previously described ^53,58,59^.

### THP-1 priming and HIV-1 infection

THP-1 cells were seeded at 1 × 10^5^ cells/well in 96-well U-bottom plates in media containing 500 ng/mL Pam3CSK4 (Invivogen) for 4-6 hours then treated with either Val-boroPro (10 μM) or nigericin (5 μg/mL) or infected with HIV-1_LAI_ or HIV-1_LAI-VSVG_ in the presence of 20 μg/mL DEAE-Dextran via spinoculation at 1100 x *g* for 30 minutes at 30°C. 24 hours post-infection or VbP treatment (two hours for nigericin), supernatants were collected for IL-1β quantification (see IL-1R reporter assay) and cells were stained with propidium iodide dye and fixed for flow cytometry.

### IL-1R reporter assay

To quantify the IL-1β secretion, we used HEK-Blue IL-1β reporter cells (Invivogen) whereby binding of IL-1β to the surface receptor IL-1R1 results in the downstream activation of NF-kB and subsequent production of secreted embryonic alkaline phosphatase (SEAP) in a dose-dependent manner as previously described ^28^. SEAP levels were detected using a colorimetric substrate assay, QUANTI-Blue (Invivogen), by measuring an increase in absorbance at OD655. Culture supernatant from treated or infected THP-1 cells was transferred to HEK-Blue IL-1β reporter cells plated in 96-well format in a total volume of 200 μL per well at 5 × 10^5^ cells/well. On the same plate, serial dilutions of recombinant human IL-1β (Peptrotech) were added to generate a standard curve for each assay. After 24 hours, SEAP levels were assayed by adding 50 μL of the supernatant from HEK-Blue IL-1β reporter cells 150 μL of QUANTI-Blue colorimetric substrate along with 0.25% Tween-20 to neutralize HIV virions in supernatant before readout. After incubation at 37°C for 15–30 minutes, absorbance at OD655 was measured on an Epoch Microplate Spectrophotometer (BioTek) and absolute levels of IL-1β were calculated relative to the standard curve.

## Supplemental information

**Table S1. List of primers, gBlocks and sgRNA sequences**

**Table S2: List of Antibodies**

## Notes

### Competing Interest Statement

The authors have declared no competing interest.

## References

1. Sharp, P.M., and Hahn, B.H. (2011). Origins of HIV and the AIDS Pandemic. Cold Spring Harb. Perspect. Med. 1, a006841. 10.1101/cshperspect.a006841.

2. Aghokeng, A.F., Bailes, E., Loul, S., Courgnaud, V., Mpoudi-Ngolle, E., Sharp, P.M., Delaporte, E., and Peeters, M. (2007). Full-length sequence analysis of SIVmus in wild populations of mustached monkeys (Cercopithecus cephus) from Cameroon provides evidence for two co-circulating SIVmus lineages. Virology 360, 407–418. 10.1016/j.virol.2006.10.048.

3. Bailes, E., Gao, F., Bibollet-Ruche, F., Courgnaud, V., Peeters, M., Marx, P.A., Hahn, B.H., and Sharp, P.M. (2003). Hybrid Origin of SIV in Chimpanzees. Science 300, 1713–1713. 10.1126/science.1080657.

4. Sharp, P.M., and Hahn, B.H. (2010). The evolution of HIV-1 and the origin of AIDS. Philos. Trans. Biol. Sci. 365, 2487–2494.

5. Keele, B.F., Jones, J.H., Terio, K.A., Estes, J.D., Rudicell, R.S., Wilson, M.L., Li, Y., Learn, G.H., Beasley, T.M., Schumacher-Stankey, J., et al. (2009). Increased mortality and AIDS-like immunopathology in wild chimpanzees infected with SIVcpz. Nature 460, 515–519. 10.1038/nature08200.

6. Paiardini, M., and Müller-Trutwin, M. (2013). HIV-associated chronic immune activation. Immunol. Rev. 254, 78–101. 10.1111/imr.12079.

7. Kaur, A., Grant, R.M., Means, R.E., McClure, H., Feinberg, M., and Johnson, R.P. (1998). Diverse Host Responses and Outcomes following Simian Immunodeficiency Virus SIVmac239 Infection in Sooty Mangabeys and Rhesus Macaques. J. Virol. 72, 9597–9611.

8. Daugherty, M.D., and Malik, H.S. (2012). Rules of Engagement: Molecular Insights from Host-Virus Arms Races. 27.

9. Parrish, C.R., Holmes, E.C., Morens, D.M., Park, E.-C., Burke, D.S., Calisher, C.H., Laughlin, C.A., Saif, L.J., and Daszak, P. (2008). Cross-Species Virus Transmission and the Emergence of New Epidemic Diseases. Microbiol. Mol. Biol. Rev. 72, 457–470. 10.1128/MMBR.00004-08.

10. Broz, P., and Dixit, V.M. (2016). Inflammasomes: mechanism of assembly, regulation and signalling. Nat. Rev. Immunol. 16, 407–420. 10.1038/nri.2016.58.

11. Harapas, C.R., Steiner, A., Davidson, S., and Masters, S.L. (2018). An Update on Autoinflammatory Diseases: Inflammasomopathies. Curr. Rheumatol. Rep. 20, 40. 10.1007/s11926-018-0750-4.

12. Taabazuing, C.Y., Griswold, A.R., and Bachovchin, D.A. (2020). The NLRP1 and CARD8 inflammasomes. Immunol. Rev. 297, 13–25. 10.1111/imr.12884.

13. D’Osualdo, A., Weichenberger, C.X., Wagner, R.N., Godzik, A., Wooley, J., and Reed, J.C. (2011). CARD8 and NLRP1 Undergo Autoproteolytic Processing through a ZU5-Like Domain. PLoS ONE 6, e27396. 10.1371/journal.pone.0027396.

14. Fink, S.L., and Cookson, B.T. (2005). Apoptosis, Pyroptosis, and Necrosis: Mechanistic Description of Dead and Dying Eukaryotic Cells. Infect. Immun. 73, 1907–1916. 10.1128/IAI.73.4.1907-1916.2005.

15. Sharif, H., Hollingsworth, L.R., Griswold, A.R., Hsiao, J.C., Wang, Q., Bachovchin, D.A., and Wu, H. (2021). Dipeptidyl peptidase 9 sets a threshold for CARD8 inflammasome formation by sequestering its active C-terminal fragment. Immunity 54, 1392-1404.e10. 10.1016/j.immuni.2021.04.024.

16. Johnson, D.C., Okondo, M.C., Orth, E.L., Rao, S.D., Huang, H.-C., Ball, D.P., and Bachovchin, D.A. (2020). DPP8/9 inhibitors activate the CARD8 inflammasome in resting lymphocytes. Cell Death Dis. 11, 628. 10.1038/s41419-020-02865-4.

17. Johnson, D.C., Taabazuing, C.Y., Okondo, M.C., Chui, A.J., Rao, S.D., Brown, F.C., Reed, C., Peguero, E., de Stanchina, E., Kentsis, A., et al. (2018). DPP8/9 inhibitor-induced pyroptosis for treatment of acute myeloid leukemia. Nat. Med. 24, 1151–1156. 10.1038/s41591-018-0082-y.

18. Rao, S.D., Chen, Q., Wang, Q., Orth-He, E.L., Saoi, M., Griswold, A.R., Bhattacharjee, A., Ball, D.P., Huang, H.-C., Chui, A.J., et al. (2022). M24B aminopeptidase inhibitors selectively activate the CARD8 inflammasome. Nat. Chem. Biol. 18, 565–574. 10.1038/s41589-021-00964-7.

19. Nadkarni, R., Chu, W.C., Lee, C.Q.E., Mohamud, Y., Yap, L., Toh, G.A., Beh, S., Lim, R., Fan, Y.M., Zhang, Y.L., et al. (2022). Viral proteases activate the CARD8 inflammasome in the human cardiovascular system. J. Exp. Med. 219, e20212117. 10.1084/jem.20212117.

20. Tsu, B.V., Agarwal, R., Gokhale, N.S., Kulsuptrakul, J., Ryan, A.P., Castro, L.K., Beierschmitt, C.M., Turcotte, E.A., Fay, E.J., Vance, R.E., et al. (2022). Host specific sensing of coronaviruses and picornaviruses by the CARD8 inflammasome. 2022.09.21.508960. 10.1101/2022.09.21.508960.

21. Wang, Q., Gao, H., Clark, K.M., Mugisha, C.S., Davis, K., Tang, J.P., Harlan, G.H., DeSelm, C.J., Presti, R.M., Kutluay, S.B., et al. (2021). CARD8 is an inflammasome sensor for HIV-1 protease activity. Science 371, eabe1707. 10.1126/science.abe1707.

22. Robinson, K.S., Teo, D.E.T., Tan, K.S., Toh, G.A., Ong, H.H., Lim, C.K., Lay, K., Au, B.V., Lew, T.S., Chu, J.J.H., et al. (2020). Enteroviral 3C protease activates the human NLRP1 inflammasome in airway epithelia. Science 370, eaay2002. 10.1126/science.aay2002.

23. Hsiao, J.C., Neugroschl, A.R., Chui, A.J., Taabazuing, C.Y., Griswold, A.R., Wang, Q., Huang, H.-C., Orth-He, E.L., Ball, D.P., Hiotis, G., et al. (2022). A ubiquitin-independent proteasome pathway controls activation of the CARD8 inflammasome. J. Biol. Chem. 298, 102032. 10.1016/j.jbc.2022.102032.

24. Sandstrom, A., Mitchell, P.S., Goers, L., Mu, E.W., Lesser, C.F., and Vance, R.E. (2019). Functional degradation: A mechanism of NLRP1 inflammasome activation by diverse pathogen enzymes. 11.

25. Barbian, H.J., Decker, J.M., Bibollet-Ruche, F., Galimidi, R.P., West, A.P., Learn, G.H., Parrish, N.F., Iyer, S.S., Li, Y., Pace, C.S., et al. (2015). Neutralization Properties of Simian Immunodeficiency Viruses Infecting Chimpanzees and Gorillas. mBio 6, e00296–15. 10.1128/mBio.00296-15.

26. Keele, B.F., Van Heuverswyn, F., Li, Y., Bailes, E., Takehisa, J., Santiago, M.L., Bibollet-Ruche, F., Chen, Y., Wain, L.V., Liegeois, F., et al. (2006). Chimpanzee Reservoirs of Pandemic and Nonpandemic HIV-1. Science 313, 523–526. 10.1126/science.1126531.

27. Tsu, B.V., Fay, E.J., Nguyen, K.T., Corley, M.R., Hosuru, B., Dominguez, V.A., and Daugherty, M.D. (2021). Running With Scissors: Evolutionary Conflicts Between Viral Proteases and the Host Immune System. Front. Immunol. 12.

28. Tsu, B.V., Beierschmitt, C., Ryan, A.P., Agarwal, R., Mitchell, P.S., and Daugherty, M.D. (2021). Diverse viral proteases activate the NLRP1 inflammasome. eLife 10, e60609. 10.7554/eLife.60609.

29. Figueiredo, A., Moore, K.L., Mak, J., Sluis-Cremer, N., Bethune, M.-P. de, and Tachedjian, G. (2006). Potent Nonnucleoside Reverse Transcriptase Inhibitors Target HIV-1 Gag-Pol. PLOS Pathog. 2, e119. 10.1371/journal.ppat.0020119.

30. Trinité, B., Zhang, H., and Levy, D.N. (2019). NNRTI-induced HIV-1 protease-mediated cytotoxicity induces rapid death of CD4 T cells during productive infection and latency reversal. Retrovirology 16, 17. 10.1186/s12977-019-0479-9.

31. Tabler, C.O., Wegman, S.J., Chen, J., Shroff, H., Alhusaini, N., and Tilton, J.C. (2022). The HIV-1 Viral Protease Is Activated during Assembly and Budding Prior to Particle Release. J. Virol. 96, e02198–21. 10.1128/jvi.02198-21.

32. Linder, A., and Hornung, V. (2022). Inflammasomes in T cells. J. Mol. Biol. 434, 167275. 10.1016/j.jmb.2021.167275.

33. Linder, A., Bauernfried, S., Cheng, Y., Albanese, M., Jung, C., Keppler, O.T., and Hornung, V. (2020). CARD8 inflammasome activation triggers pyroptosis in human T cells. EMBO J. 39, e105071. 10.15252/embj.2020105071.

34. Ball, D.P., Taabazuing, C.Y., Griswold, A.R., Orth, E.L., Rao, S.D., Kotliar, I.B., Vostal, L.E., Johnson, D.C., and Bachovchin, D.A. (2020). Caspase-1 interdomain linker cleavage is required for pyroptosis. Life Sci. Alliance 3. 10.26508/lsa.202000664.

35. McKee, C.M., and Coll, R.C. (2020). NLRP3 inflammasome priming: A riddle wrapped in a mystery inside an enigma. J. Leukoc. Biol. 108, 937–952. 10.1002/JLB.3MR0720-513R.

36. Álvarez, E., Castelló, A., Menéndez-Arias, L., and Carrasco, L. (2006). HIV protease cleaves poly(A)-binding protein. Biochem. J. 396, 219–226. 10.1042/BJ20060108.

37. Chahroudi, A., Bosinger, S.E., Vanderford, T.H., Paiardini, M., and Silvestri, G. (2012). Natural SIV Hosts: Showing AIDS the Door. Science 335, 1188–1193. 10.1126/science.1217550.

38. Doitsh, G., Galloway, N.L., Geng, X., Yang, Z., Monroe, K.M., Zepeda, O., Hunt, P.W., Hatano, H., Sowinski, S., Muñoz-Arias, I., et al. (2014). Pyroptosis drives CD4 T-cell depletion in HIV-1 infection. Nature 505, 509–514. 10.1038/nature12940.

39. Muema, D.M., Akilimali, N.A., Ndumnego, O.C., Rasehlo, S.S., Durgiah, R., Ojwach, D.B.A., Ismail, N., Dong, M., Moodley, A., Dong, K.L., et al. (2020). Association between the cytokine storm, immune cell dynamics, and viral replicative capacity in hyperacute HIV infection. BMC Med. 18, 81. 10.1186/s12916-020-01529-6.

40. Arditi, M., Kabat, W., and Yogev, R. (1991). Serum tumor necrosis factor alpha, interleukin 1-beta, p24 antigen concentrations and CD4+ cells at various stages of human immunodeficiency virus 1 infection in children. Pediatr. Infect. Dis. J. 10, 450–454.

41. Monroe, K.M., Yang, Z., Johnson, J.R., Geng, X., Doitsh, G., Krogan, N.J., and Greene, W.C. (2014). IFI16 DNA Sensor Is Required for Death of Lymphoid CD4 T-cells Abortively Infected with HIV. Science 343, 428–432. 10.1126/science.1243640.

42. Zhang, C., Song, J.-W., Huang, H.-H., Fan, X., Huang, L., Deng, J.-N., Tu, B., Wang, K., Li, J., Zhou, M.-J., et al. (2021). NLRP3 inflammasome induces CD4+ T cell loss in chronically HIV-1–infected patients. J. Clin. Invest. 131. 10.1172/JCI138861.

43. Muñoz-Arias, I., Doitsh, G., Yang, Z., Sowinski, S., Ruelas, D., and Greene, W.C. (2015). Blood-Derived CD4 T Cells Naturally Resist Pyroptosis During Abortive HIV-1 Infection. Cell Host Microbe 18, 463–470. 10.1016/j.chom.2015.09.010.

44. Sandler, N.G., and Douek, D.C. (2012). Microbial translocation in HIV infection: causes, consequences and treatment opportunities. Nat. Rev. Microbiol. 10, 655–666. 10.1038/nrmicro2848.

45. Kim, J.G., and Shan, L. (2022). Beyond Inhibition: A Novel Strategy of Targeting HIV-1 Protease to Eliminate Viral Reservoirs. Viruses 14, 1179. 10.3390/v14061179.

46. Moore, K.P., Schwaid, A.G., Tudor, M., Park, S., Beshore, D.C., Converso, A., Shipe, W.D., Anand, R., Lan, P., Moningka, R., et al. (2022). A Phenotypic Screen Identifies Potent DPP9 Inhibitors Capable of Killing HIV-1 Infected Cells. ACS Chem. Biol. 17, 2595–2604. 10.1021/acschembio.2c00515.

47. Chung, Y., Chang, S.H., Martinez, G.J., Yang, X.O., Nurieva, R., Kang, H.S., Ma, L., Watowich, S.S., Jetten, A.M., Tian, Q., et al. (2009). Critical Regulation of Early Th17 Cell Differentiation by Interleukin-1 Signaling. Immunity 30, 576–587. 10.1016/j.immuni.2009.02.007.

48. Rider, P., Carmi, Y., Guttman, O., Braiman, A., Cohen, I., Voronov, E., White, M.R., Dinarello, C.A., and Apte, R.N. (2011). IL-1α and IL-1β Recruit Different Myeloid Cells and Promote Different Stages of Sterile Inflammation. J. Immunol. 187, 4835–4843. 10.4049/jimmunol.1102048.

49. Min, A.K., Fortune, T., Rodriguez, N., Hedge, E., and Swartz, T.H. (2022). Inflammasomes as mediators of inflammation in HIV-1 infection. Transl. Res. 10.1016/j.trsl.2022.07.008.

50. Busnadiego, I., Kane, M., Rihn, S.J., Preugschas, H.F., Hughes, J., Blanco-Melo, D., Strouvelle, V.P., Zang, T.M., Willett, B.J., Boutell, C., et al. (2014). Host and Viral Determinants of Mx2 Antiretroviral Activity. J. Virol. 88, 7738–7752. 10.1128/JVI.00214-14.

51. Poss, M., and Overbaugh, J. (1999). Variants from the Diverse Virus Population Identified at Seroconversion of a Clade A Human Immunodeficiency Virus Type 1-Infected Woman Have Distinct Biological Properties. J. Virol. 73, 5255–5264.

52. Guyader, M., Emerman, M., Sonigo, P., Clavel, F., Montagnier, L., and Alizon, M. (1987). Genome organization and transactivation of the human immunodeficiency virus type 2. Nature 326, 662–669. 10.1038/326662a0.

53. Peden, K., Emerman, M., and Montagnier, L. (1991). Changes in growth properties on passage in tissue culture of viruses derived from infectious molecular clones of HIV-1LAI, HIV-1MAL, and HIV-1ELI. Virology 185, 661–672. 10.1016/0042-6822(91)90537-L.

54. Labun, K., Montague, T.G., Krause, M., Torres Cleuren, Y.N., Tjeldnes, H., and Valen, E. (2019). CHOPCHOP v3: expanding the CRISPR web toolbox beyond genome editing. Nucleic Acids Res. 47, W171–W174. 10.1093/nar/gkz365.

55. Schmid-Burgk, J.L., Schmidt, T., Gaidt, M.M., Pelka, K., Latz, E., Ebert, T.S., and Hornung, V. (2014). OutKnocker: a web tool for rapid and simple genotyping of designer nuclease edited cell lines. Genome Res. 24, 1719–1723. 10.1101/gr.176701.114.

56. Schmidt, T., Schmid-Burgk, J.L., Ebert, T.S., Gaidt, M.M., and Hornung, V. (2016). Designer Nuclease-Mediated Generation of Knockout THP1 Cells. In TALENs Methods in Molecular Biology., R. Kühn, W. Wurst, and B. Wefers, eds. (Springer New York), pp. 261–272. 10.1007/978-1-4939-2932-0_19.

57. OhAinle, M., Helms, L., Vermeire, J., Roesch, F., Humes, D., Basom, R., Delrow, J.J., Overbaugh, J., and Emerman, M. (2018). A virus-packageable CRISPR screen identifies host factors mediating interferon inhibition of HIV. eLife 7, e39823. 10.7554/eLife.39823.

58. Bartz, S.R., and Vodicka, M.A. (1997). Production of High-Titer Human Immunodeficiency Virus Type 1 Pseudotyped with Vesicular Stomatitis Virus Glycoprotein. Methods 12, 337–342. 10.1006/meth.1997.0487.

59. Gummuluru, S., Rogel, M., Stamatatos, L., and Emerman, M. (2003). Binding of Human Immunodeficiency Virus Type 1 to Immature Dendritic Cells Can Occur Independently of DC-SIGN and Mannose Binding C-Type Lectin Receptors via a Cholesterol-Dependent Pathway. J. Virol. 77, 12865–12874. 10.1128/JVI.77.23.12865-12874.2003.

