## Supporting Information for "A human-specific motif facilitates CARD8 inflammasome activation after HIV-1 infection"

### Supplemental Tables

**Table S1: List of primers, gBlocks and sgRNA sequences**

| Type | Target | Name | Sequence | Description |
|---|---|---|---|---|
| primer | human CARD8 | pcDNA3.1_BamHI_mCherry_F | TTGGTACCGAGCTCGGATCCATGGTGAGCAAGGGC | forward primer to clone mC-CARD8 into pcDNA3.1 backbone |
| primer | human CARD8 | pcDNA3.1_EcoRI_CARD8_Cterm_R | GCTGGATATCTGCAGAATTCTTACAAATTCTGCTGTCTAAGATAG | reverse primer to clone mC-CARD8 into pcDNA3.1 backbone |
| primer | human CARD8 | CARD8_cpz_F60L | GGAATTTTTCTTCAGGCTGAGGC | forward primer for T>C mutation in CARD8 introducing a phenylalanine to leucine mutation |
| primer | human CARD8 | CARD8_cpz_F60L | AGCCTGAAGAAAAATTCCAGTTTTTGTGTAT | reverse primer for A>G mutation in CARD8 introducing a phenylalanine to leucine mutation |
| primer | human CARD8 | CARD8_OWM_F60S | GGAATTTTTCTCAGGCTGAGGC | forward primer for T>C mutation in CARD8 introducing a phenylalanine to serine mutation |
| primer | human CARD8 | CARD8_OWM_F60S | AGCCTGAGAAAAAATTCCAGTTTTTGTGTAT | reverse primer for A>G mutation in CARD8 introducing a phenylalanine to serine mutation |
| primer | human CARD8 | CARD8_F60A_internal_F | GGAATTTTTGCGCAGGCTGAGGC | forward primer for TTT>GCG mutation in CARD8 introducing a phenylalanine to alanine mutation |
| primer | human CARD8 | CARD8_F60A_internal_R | AGCCTGCGCAAAAATTCCAGTTTTTGTGTAT | reverse primer for AAA>CGC mutation in CARD8 introducing a phenylalanine to alanine mutation |
| primer | mCherry | mCherry_F | ATGGTGAGCAAGGGCGAG | forward primer to amplify mCherry-CARD8 |
| primer | human CARD8 | CARD8_Cterm_R | TTACAAATTCTGCTGTCTAAGATAGGACAC | reverse primer to amplify CARD8 |
| primer | human CARD8 | CARD8_seq_internal_F | ATGGAAAAAAAGGAGTGTCC | forward primer for sequence verifying CARD8 construct |

**Table S1 (continued): List of primers, gBlocks and sgRNA sequences**

| Type | Target | Name | Sequence | Description |
|---|---|---|---|---|
| primer | human CARD8 | CARD8_Vance_geno_F | GATGTTGCAGTGAGCCAAGA | forward primer for verifying CARD8 KO at sgRNA site |
| primer | human CARD8 | CARD8_Vance_geno_R | CGTCTCACTGCTGTTGTGGT | reverse primer for verifying CARD8 KO at sgRNA site |
| primer | human CARD8 | CARD8_LKO_SfiI_Kozak_F | AAAAGGCCGAGAGGGCCGAATTCGCCACCATGGAAAAAAAGGAGTGTCC | forward primer for cloning CARD8 into pLKO vector with Kozak sequence |
| primer | human CARD8 | CARD8_LKO_linker_Myc_SfiI_R | AAAAGGCCAGAGAGGCCCTAGAGATCCTCTTCTGAGATGAGTTTTTGTTCGCTAGCTGCCGCTCCGCTTCCCAAATTCTGCTGTCTAAGATAGGACAC | reverse primer for cloning CARD8 into pLKO vector with Myc tag |
| primer | chimpanzee CARD8 | NotI_linker_cpz_CARD8_F | AAAAGCGGCCGCAATGGAAAAAAAGGAGTTTCC | forward primer for cloning chimpanzee CARD8 into pcDNA3.1 vector |
| primer | chimpanzee CARD8 | CARD8_cterm_NotI_pcDNA3_R | CTCTAGACTCGAGCGGCCGCCACTGTGCTGGATATCTGCAGAATTCTTACAAATTCTGCT | reverse primer for cloning chimpanzee CARD8 into pcDNA3.1 vector |
| primer | chimpanzee CARD8 | CARD8_LKO_SfiI_Kozak_cpz_CARD8_F | AAAAGGCCGAGAGGGCCGAATTCGCCACCATGGAAAAAAAGGAGTTTCC | forward primer for cloning CARD8 into pLKO vector with Kozak sequence |
| primer | CARD8 | pLKO_check_F1 | GGAGGTCTATATAAGCAGAGCTCTCCC | forward primer to amplify insert in pLKO vector |
| primer | CARD8 | pLKO_check_R1 | CTACTATTCTTTCCCCTGCACTGTACCC | reverse primer to amplify insert in pLKO vector |
| primer | CARD8 | pLKO_check_F2 | CAGTGATAGAGATCTCCCTATCAG | forward primer to sequence insert in pLKO vector |
| primer | CARD8 | pLKO_check_R2 | GGATGAATACTGCCATTTGTCTCGAGG | reverse primer to amplify sequence in pLKO vector |
| gBlock | chimpanzee CARD8 | chimp_CARD8 | see NCBI accession XM_024351500.1 | gblock for chimpanzee CARD8 |
| sgRNA | CARD8 | CARD8_sgRNA | TTGTTAGCAAGGCGTCGCTGGGG | sgRNA for generating CARD8 KO |

**Table S2: List of Antibodies**

| Product | Catalog Number | Clone | Dilution Ratio | Purpose | Company | City | Country | State |
|---|---|---|---|---|---|---|---|---|
| mCherry mouse mAb | 632543 | – | 1:1000 | Western blot | Takara Bio | Kusatsu | Japan | – |
| p24 mouse mAb | ARP-3537 | – | 1:5000 | Western blot | NIH HIV Reagents Program | Manassas | USA | Virginia |
| vinculin mouse mAb | sc-73614 | 7F9 | 1:5000 | Western blot | Santa Cruz Biosciences | Santa Cruz | USA | California |
| CARD8 Rabbit pAb (C-terminal) | ab24186 | – | 1:1000 | Western blot | Abcam | Cambridge | United Kingdom | – |
| HIV-1 core antigen-FITC | 6604665 | KC57 | 1:300 | Flow cytometry | Beckman Coulter | Indianapolis | USA | Indiana |
